# Cross-feeding shapes both competition and cooperation in microbial ecosystems

**DOI:** 10.1101/2021.10.10.463852

**Authors:** Pankaj Mehta, Robert Marsland

**Author notes:** Pontifical University of the Holy Cross, Rome, Italy (Robert Marsland).

## Abstract

Recent work suggests that cross-feeding – the secretion and consumption of metabolic biproducts by microbes – is essential for understanding microbial ecology. Yet how cross-feeding and competition combine to give rise to ecosystem-level properties remains poorly understood. To address this question, we analytically analyze the Microbial Consumer Resource Model (MiCRM), a prominent ecological model commonly used to study microbial communities. Our mean-field solution exploits the fact that unlike replicas, the cavity method does not require the existence of a Lyapunov function. We use our solution to derive new species-packing bounds for diverse ecosystems in the presence of cross-feeding, as well as simple expressions for species richness and the abundance of secreted resources as a function of cross-feeding (metabolic leakage) and competition. Our results show how a complex interplay between competition for resources and cooperation resulting from metabolic exchange combine to shape the properties of microbial ecosystems.

---

Microbial communities are found everywhere on the globe from hot springs to human bodies [1, 2]. Large-scale surveys suggest that a these microbial communities are extremely diverse, with hundreds to thousands of distinct microbial species all coexisting in a single environment. Recent lab experiments suggest that that diverse communities can also stably coexist even in simple environments with just a handful of externally supplied resources suggesting that classical ecological theories of competition must be modified to describe microbial ecosystems [3–7].

It is now believed that metabolic interactions facilitated by cross-feeding – the secretion and consumption of metabolic biproducts by microbes – play a fundamental role in shaping the structure and function of microbial ecosystems. [3, 6, 8–11].This has led to a substantial body of work trying to extend classical ecological model to incorporate cross-feeding interactions [3, 6, 12–15]. One prominent example of this is the Microbial Consumer Resource Model (MiCRM) which extends classical ecological consumer resource models [16–20] by including cross-feeding interactions between consumers [13, 21, 22]. The MiCRM has been successfully used to explain both laboratory experiments and the natural patterns found in large-scale surveys [5, 21–23].

Whereas classical consumer resource models emphasize the importance of competition in shaping ecosystems [18–20], the MiCRM includes both competition for resources as well as cooperation between microbes facilitated by cross-feeding interactions. A fundamental question in microbial ecology is to understand how this interplay between competition and cooperation shapes the properties of diverse communities [11]. Answering this question has proven difficult due to a lack of analytic results. The MiCRM, in contrast with many other ecological models, cannot be cast as an optimization problem [16, 17, 24, 25] and for this reason is not amenable to analysis using the replica method [26–28]. Here, we use the cavity method, another technique from the statistical physics of disordered systems [29–31], to derive a mean-field theory for the steady-state solutions for the MiCRM. Unlike replicas, the cavity method does not require the existence of an optimization functional making it ideally suited for analyzing the MiCRM [32]. Since this calculation is quite technically involved, we limit ourselves in the main text to presenting the results and implications. A detailed derivation is provided in the appendix.

## Model

We briefly summarize Microbial consumer resource model (MiCRM) (see [13, 21] and appendix for details). Species *i* = 1… *S* with abundances *N_i_* can consume *M* potential resources *R_β_* (*β* = 1… *M*) that can exist in the environment. A fraction (1 – *l*) of the energy in consumed resources is used for growth while the remaining energy fraction 0 ≤ *l* ≤ 1 is released back into the environment as metabolic biproducts (see Fig. 1). This is described by the equation:

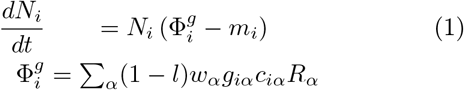

where *w_α_* is the value of one unit of resource to a species (e.g. ATPs that can be extracted); *c_iα_* is the rate at which species *i* consumes resource *α*, *m_i_* is the minimum amount of resources that must be consumed in order to have a positive growth rate, and *g_iα_* is a factor that converts consumption into a growth rate for species *i* when consuming resource *α*. To derive our analytic solution, we consider the case when the consumer preferences *c_iα_* are drawn randomly with 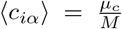 and 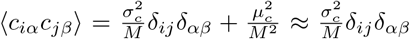. We have numerically checked that in the thermodynamic limit where *M*, *S* → ∞ with *γ* = *M*/*S* fixed that our analytic expressions hold even when *c_iα_* are drawn from distributions where the *c_iα_* are strictly positive (see Figures S1 and 2).

**FIG. 1.**
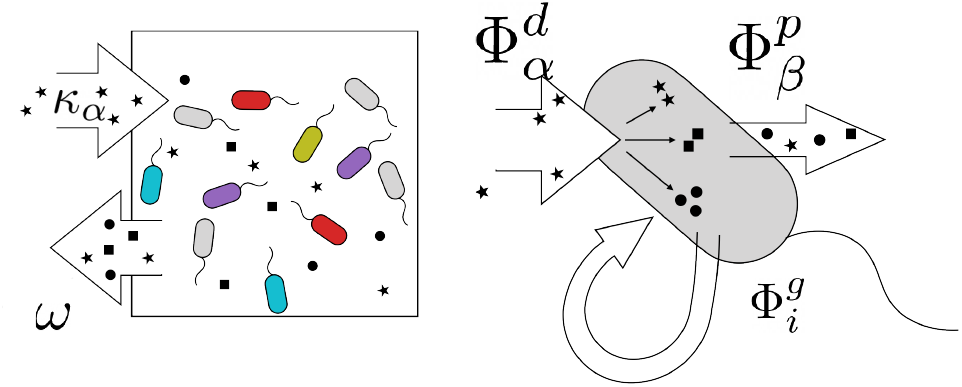
Microbial Consumer Resource Model (Mi-CRM). Resources {*R_α_*} are supplied externally into the environment at a rate *κ_α_* and degraded at a rate *ω*. Microbes (also called consumers) with abundances {*N_i_*} consume resources from the environment 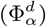, using some of the energy to grow 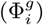, while the remaining energy is secreted back into the environment as metabolic biproducts 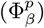. We assume there is a single externally supplied resource (*α* = *E*) so that *κ_α_* =0 if *α* ≠ *E*.

In the MiCRM, the resources *R_α_* satisfy their own dy namical equations:

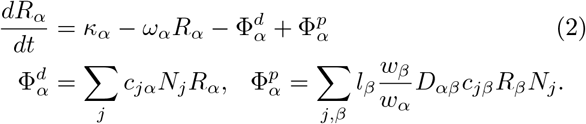

The first two terms on the right hand side of the top equation describe the dynamics of *R_α_* in the absence of microbes, 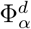 describes resource consumption, and 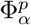 the production of resources due to cross feeding. The cross-feeding matrix *D_αβ_* encodes the fraction of energy leaked from resource *β* that goes into resource *α*. Motivated by the universality of metabolism, we assume that the *D_αβ_* is the same for all species *i* and each row can be described using a Dirichlet distribution. To ensure a good thermodynamic limit, we further assume that elements of *D_αβ_* for a fixed *β* follow a Dirichlet distribution with shape parameters {*α_j_*} which scale like 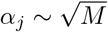. Under this assumption, to leading order in *M* we can ignore the anti-correlations between the elements of *D_αβ_* when deriving our mean-field equations and instead approximate the the elements of *D_αβ_* as independent, normal random variables with 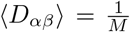 and 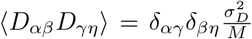 (see appendix). We emphasize that our numerical simulations simulate the full dynamics and do not rely on this approximation.

In what follows we set *w_α_* = *w* and *g_iα_* = *g* for all *α* and *i*. We also assume that the *m_i_* are independent normal variables with mean *m* and variance 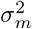. Finally, inspired by recent experiments [3–7] we focus on the case where just one of the resources *R_E_* is supplied externally at a rate 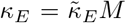, and all the other resources result from cross-feeding (i.e *κ_α_* = 0 for *α* ≠ *E*). This corresponds to the high-energy regime that supports diverse communities in [13].

## Parameters, competition, and cross-feeding

We are especially interested in understanding the interplay between competition and cooperation. In all consumer resource models, consumers/microbes that have similar preference for resources compete more than those who have very different consumer preferences since they occupy similar metabolic niches. This intuition can be generalized to diverse communities by looking at the quantity *σ_c_*/*μ_c_* which characterizes the diversity of consumer resource preferences in the regional species pool. When *σ_c_* ≪ *μ_c_*, all species have the similar consumer preferences and there is lots of competition, whereas when *σ_c_* ≫ *μ_c_* there is much less competition between species since species have distinct consumer preferences. For this reason, we interpret the ratio *σ_c_*/*μ_c_* as a proxy for (the inverse of) the amount of competition in the regional species pool.

The amount of cross-feeding in the community is controlled by the leakage parameter *l* which governs the fraction of energy consumed by microbes that is released as metabolic biproducts. When *l* = 0, all the energy microbes consume is used for growth and Φ_*p*_ = 0. In this case, our equations for the MiCRM reduce to the usual equations for a consumer resource model without cross-feeding [26, 33]. In contrast, if *l* = 1 all the energy is released as metatabolic biproducts and species cannot grow. More generally, increasing *l* increases the fraction of energy released as metabolic biproducts and hence serves as a proxy for the amount of cross-feeding in the community [13].

## Mean-field equations

In the appendix, we derive mean-field equations for the MiCRM using the cavity method. These calculations are extremely long and technical and here we confine ourselves to simply stating the final results. As in the consumer resource model with linear resource dynamics [33], to derive our expressions we assume replica symmetry and solve the cavity-equations using a Taylor expansion that approximates the fluxes 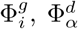, and 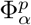 using their means and variances. We note that the the expectation value 〈*X*〉 denotes averages over both random realizations of the parameters (quenched averages) as well as species/resource abundance distributions. The end result of the cavity procedure is a set of self-consistency equations relating the fraction of surviving species *ϕ_N_*, the mean 〈*N*〉 and second-moment 〈*N*^2^〉 of the species abundance distribution, the mean 〈*R*〉 and second moment 〈*R*^2^〉 of the resources secreted through cross-feeding, and the cavity susceptibilities *χ* and *ν* (see Appendix for full definitions).

These equations can be most succinctly written in terms of the average effective degradation rate of resources in the environment 〈*ω*^eff^〉 = *ω* + *μ_c_* 〈*N*〉, which is the sum of the dilution rate *ω* and the average rate at which microbes consume resources *μ_c_*〈*N*〉. In addition, we define the parameters

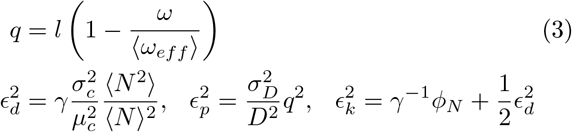

which characterize fluctuations in the fluxes appearing in Eq. 3. In terms of these quantities, the species distributions satisfy the self-consistency equations

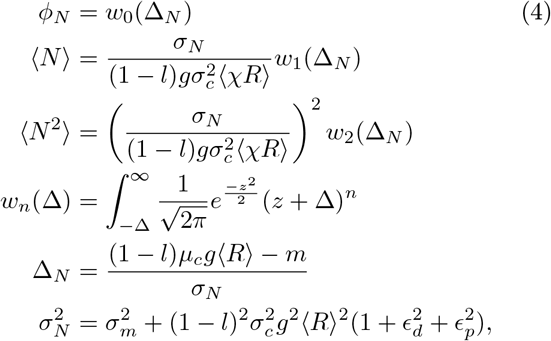

The full species abundace distribution can be calculated by noting that probability of observing a species with abundance *N*_0_ in the microbial community is given by a truncated Gaussian described of the form

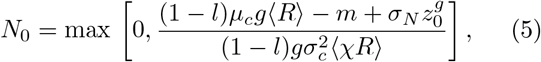

where *z_g_* is a Gaussian random normal variable describing fluctuations in the growth flux 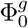 and the equation for the expectation 〈*χR*〉 is given below.

We also find that the moments of the resource abundances satisfy the self-consistency equations

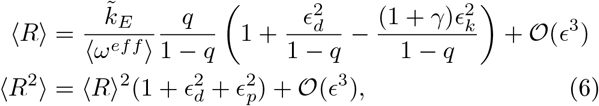

where 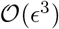 indicates that we have only kept terms to second order in *ϵ_d_*, *ϵ_p_*, and *ϵ_k_*. The full resource distribution is given by

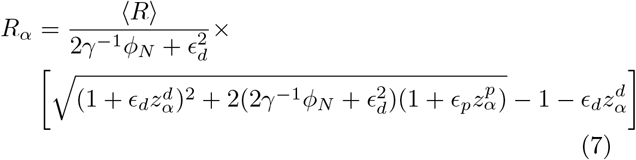

where 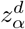 and 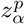 are standard random normal variables with mean zero and variance 1 describing the fluctuations in the depletion and production fluxes.

Finally, these equations must be supplemented by equations for expectations involving the cavity suscep tibilities of the form

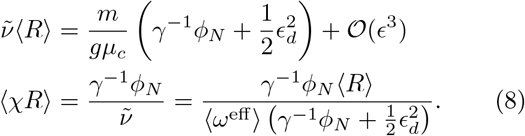

To check our expressions, we performed numerical simulations of the MiCRM using the Community Simulator package (see appendix and associated Github repository) [21]. In these numerical simulations a single externally supplied resource is supplied to the environment, but microbes can produce M=300 additional resources through cross-feeding (leakage rate *l* = 0.8). The consumer preference matrix *c_iα_* were chose from a binomial distribution. Figure S1 shows that the analytic expressions given by Eqs. 5 and 7 for the species and resource distributions (solid lines) agree well with distributions obtained numerically suggesting the cavity equations are capturing the essential ecology of the MiCRM.

## Average metabolite abundance depends is controlled by leakage

One striking result of our analytic expressions is that to leading order in *ϵ* = {*ϵ_d_*, *ϵ_p_*, *ϵ_k_*} the average abundance of metabolic biproducts 〈*R*〉 depends only on the leakage rate *l* and the effective degradation rate 〈*ω_eff_*〉 but not on the amount of competition in the regional species pool as measured by *σ_c_*/*μ_c_* (see Eq. 6). When 〈*ω_eff_*〉 » 〈*ω*〉, we can use energy conservation to derive a particularly simple approximate expression for the average metabolite abundances

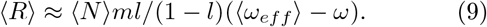

A comparison of this expression with numerics is shown in Fig. 2a. From this, we conclude that metabolite concentrations are affected by the regional species pool only through the average species abundance 〈*N*〉 and average resource degradation rate 〈*ω_eff_*〉. This shows the resource abundances reflect the functional structure of the community rather than taxonomy (i.e. exactly what microbes are present).

**FIG. 2.**
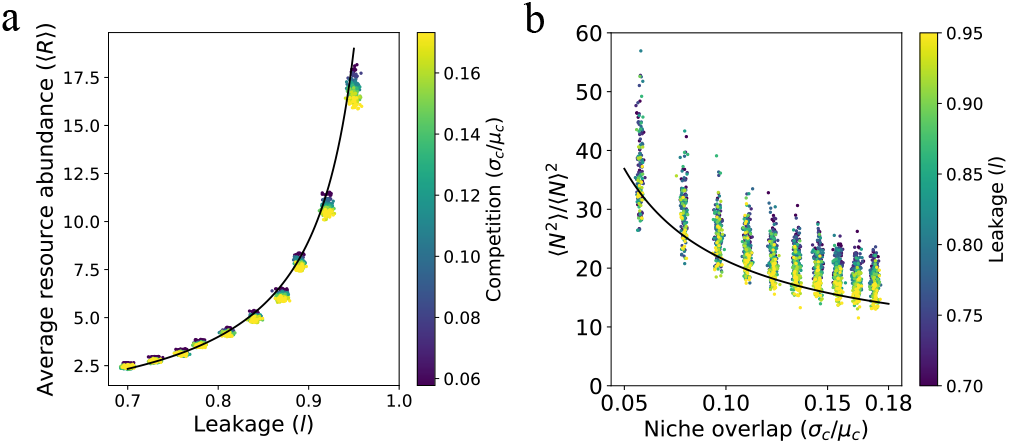
Resources and species abundances are controlled by different processes. (a) Numerical simulations of average resource abundances as a function of leakage rate *l* (x-axis) and *σ_c_*/*μ_c_* (color bar). Solid line is analytic expression from Eq. 9. (b) Numerical simulations of species 〈*N*^2^〉/〈*N*〉^2^ as a function of *σ_c_*/*μ_c_* (x-axis) and leakage rate *l* (color bar). Solid line is analytic expression given by Eq. 10. (see Fig. S1 for further numerical checks and appendix for details and parameters).

## Species distribution shaped by competition in regional species pool

The cavity solution also shows that the species abundance distribution is shaped primarily by competition (*σ_c_*/*μ_c_*) and depends much more weakly on the amount of cross-feeding as measured by the leakage rate *l*. As shown in the appendix, we find that to leading order in *ϵ* = {*ϵ_d_*, *ϵ_p_*, *ϵ_k_*}, that the quantity Δ_*N*_ depends only on 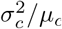. Since Δ_*N*_ is the primary determinant of species abundances, our analysis implies that both the fraction of surviving species *ϕ_N_* as well as the ratio

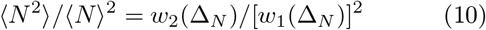

should depend strongly on the amount of competition *σ_c_*/*μ_c_* but be largely agnostic to *l*. Fig.2b shows numerical simulations confirming that this is indeed the case.

## Species packing bound

One final consequence of our analytic solution is that it allows us to derive a species packing bound for microbial ecosystems [16, 17]. In particular, we can ask how many species can co-exist in a microbial ecosystem with cross-feeding? We show in the appendix using a very similar argument to [33] that the ratio of surviving species *S*\* to metabolic biproducts *M* must be less than half (i.e *S*\*/*M* ≤ 1/2) in the thermodynamic limit. Fig. 3 show numerical simulations confirming this result. These results indicate that the species packing bound found for linear resource dynamics without cross-feeding in [33] also holds for the MiCRM where resources are primarily produced by cross-feeding.

**FIG. 3.**
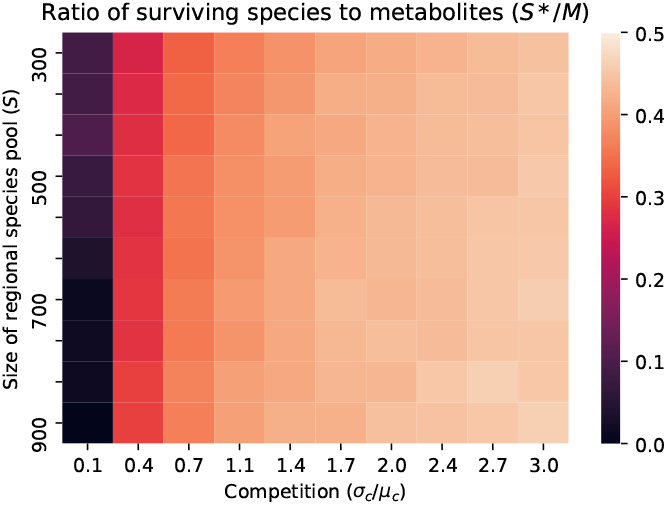
Species packing bound. Ratio of surviving species to resources (including metabolic biproducts) *S*\* /*M* for different sizes of regional species pools *S* and different choices of *σ_c_*/*μ_c_*. Notice that *S*\*/*M* ≤ 1/2, consistent analytic bound derived in appendix. Further numeric checks in Fig. S2. Parameters in appendix.

## Discussion

In this paper, we have used the cavity method to analytically calculate self-consistent meanfield equations for the MiCRM. Our solution illustrates that an intricate interplay between competition for resources and cooperation resulting from metabolic exchange shape the properties of microbial ecosystems. We found that metabolite abundances are primarily controlled by the leakage and community-level consumption rates of resources. This finding is consistent with recent experimental and theoretical work suggesting that community-level functional structure is much more conserved in similar environments than taxonomy [34, 35]. Indeed, our analytic solutions suggest that the metabolite abundances are largely agnostic to which microbes are producing or consuming resources as long as the community is sufficiently diverse.

In contrast, we find that the species abundance distribution depends primarily on the amount of competition in the regional species pool and is largely agnostic to whether metabolites were supplied externally or internally generated by the community through crossfeeding. This suggests that competition is the primary force shaping taxonomic structure. However, we emphasize that cross-feeding interactions modify the environment by creating new metabolic niches and thus fundamentally shape the terms on which microbes comepete. This basic picture is given further support by our calculation showing that species packing in the MiCRM is analogous to the bound derived for ecosystems without cross-feeding where all resources are supplied externally: the number of surviving species is always less than equal to the number of metabolites in the ecosystem. In other words, species packing in diverse communities is agnostic to how resources are generated.

There are number of promising directions worth pursuing further. In our calculations, both the consumer preferences and cross-feeding matrix were unstructured and we focused on steady-states. It should be possible to extend the calculations presented here to include taxonomic and metabolic structure in the consumption and cross-feeding matrices, as well as metabolic constraints [36–38]. More ambitiously, if will be interesting to extend our calculation to derive dynamical mean-field equations [32, 39]. Another promising direction is to understand how space and migration modify the intuitions found here [40–42]. Finally, we would like to explore if we can exploit the mean-field equations derived here to understand eco-evolutionary dynamics in the presence of cross-feeding [43]. Recent simulations and analytic arguments suggest that such eco-evolutionary dynamics may depend largely on the functional structure of the community, opening a potential avenue of extending these calculations to include evolutionary dynamics [22, 43–45].

## Acknowledgments

We thank Wenping Cui, Jason Rocks, and Josh Goldford for useful discussions. This work was supported by NIH NIGMS 1R35GM119461 and a Simons Investigator in the Mathematical Modeling of Living Systems (MMLS) award to PM. We also acknowledge support from the SSC computing cluster at BU. P.M. and R.M. contributed equally to this work.

## Supplemental Appendix

### Basic Setup

We start by briefly summarize Microbial consumer resource model (MiCRM) introduced and developed in [3, 13, 21, 23]. Species, indexed *i* = 1… *S*, grow by consuming *M* potential resources, *R_β_* (*β* = 1… *M*), that can be found in the the environment. In the MiCRM, the consumption of resources is never one-hundred percent efficient and a fraction *l_α_* of the energy in the consumed resources “leak” back into the environment as metabolic biproducts.

In this MiCRM, this is described by the equation:

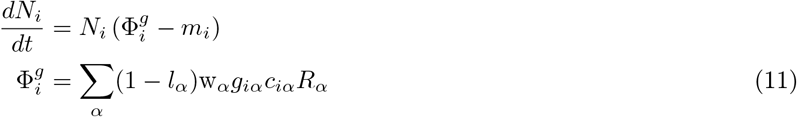

where w_*α*_ is the value of one unit of resource to a species (e.g. ATPs that can be extracted); *c_iα_* is the rate at which species *i* consumes resource *α*, *m_i_* is the minimum amount of resources that must be consumed in order to have a positive growth rate. Finally, *g_iα_* is a factor that converts consumption into a growth rate for species *i* when consuming resource *α*.

In the MiCRM, resources satisfy their own dynamical equations

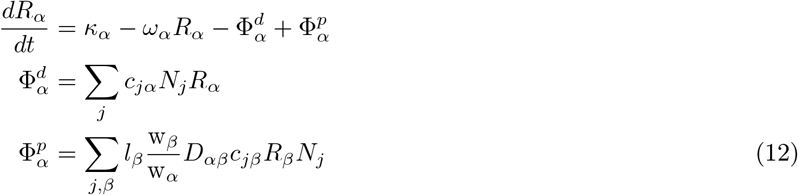

where the first two terms describe resource dynamics in the absence of microbes, 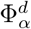, describes the depletion of resource *α* due to consumption, and the production of resources due to cross feeding from metabolic biproducts is encoded in 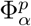. The matrix *D_αβ_* appearing in 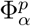 encodes the fraction of the energy released in metabolic biproduct *α* when a microbe consumes a unit of resource *β* (see below for more details). We note that these equations reduce to the equations for the original Consumer Resource Model of Levins and MacArthur in the absence of leakage (i.e. *l_α_* = 0)[16–18].

To proceed, it is helpful to introduce some additional notation and make some simplifications:

- Define the ratio of potential resources to species by

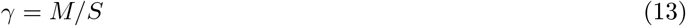
- For simplicity assume that both the growth rates and resource weights can be assumed to be constant *g_iα_* = *g* and w_*α*_ = w = 1. This is equivalent to the physical assumption that *all metabolic pathways are basically equally efficient*.
- We will also assume that all resources are leaked at the same rate so that *l_α_* = *l* and that the resource dilution rates are all equal with *ω_α_* = *ω*.
- We focus on the case where just one resource *R_E_* is supplied externally, with influx *κ_E_*, and all the others are internally generated through biproduct secretion. In other words, *κ_α_* =0 for *α* ≠ *E*.
- Finally, all our calculations will be performed in the thermodynamic limit with *M*, *S* → ∞ and the ratio *γ* fixed. For this reason, we will only keep terms that are order 1 in both *M* and *S*.
- We assume that there is no family structure in either the species consumer preferences or the cross-feeding matrix. Even this simple case has been shown to describe many real experiments on microbial communities [23].

We will consider the case when the consumer preferences *c_iα_* are drawn randomly with

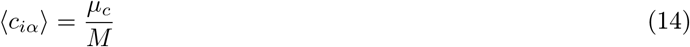

and

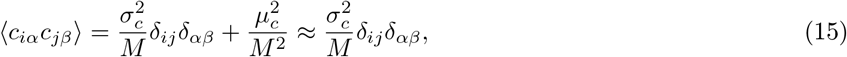

where in writing the second line we have ignored terms that are higher order in *M*. Since there are no correlations between the consumer preferences for different species, we have made the ecological assumption that *there is no family structure* in the regional species pool.

We also assume no mebaolic structure in the cross feeding matrix *D_αβ_*. In general, this matrix encodes the fraction of energy leaked from resource *β* that goes into resource *α*. Following earlier works, we assume that each row in the matrix follows a Dirichlet distribution. To ensure a good thermodynamic limit, we assume that the shape parameters {*α_j_*} of the Dirichlet distribution scale like 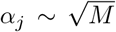 and that the sum of the shape parameters 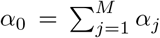 scales like *M*^3/2^. With these assumptions, the expectation, variance, and covariance of a variables {*X_i_*} draw from a Dirichlet distribution scale like

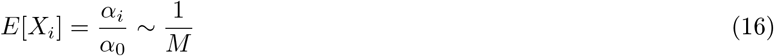

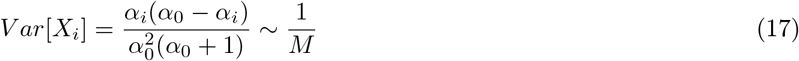

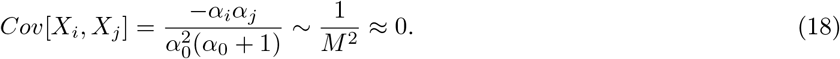

Thus, to leading order in *M* we can ignore the anti-correlations between elements of the Dirichlet distribution in the thermodynamic limit. We can translate these observation to the cross-feeding matrix *D_αβ_* to get

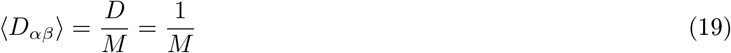

and

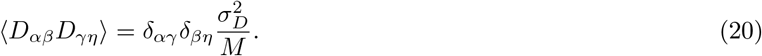

Importantly, we can ignore correlations between the elements of *D_αβ_*. Note also that by assumption *D* = 1, since *D_αβ_* is defined as a matrix that partitions a given total quantity of output flux over possible metabolites, with Σ_*α*_ *D_αβ_* = 1.

### Energy Conservation

One important feature of the MiCRM is that it explicitly accounts for energy conservation. The total power (energy per unit time) that flows into the system is just

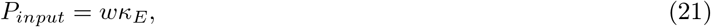

where *κ_E_* is the rate at which units of the external resource is added to the system and *w* is the energy contained in each unit of resource. At steady-state, this energy is either consumed by the microbes at a rate *P_microbe_* or lost to the environment *P_environment_* due to resources disappearing at a rate *ω_α_* (i.e. we are considering an open system where resource *α* is lost at a rate *ω_α_*). We know that by construction

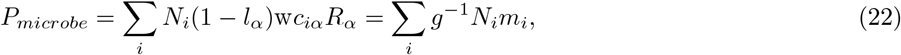

where in going to the second line we have Eq. 11, noting that at steady-state 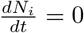 for all microbes. We also have by definition that

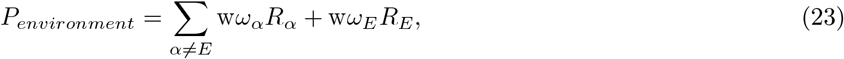

where for future convenience we have explicitly separated the externally supplied resource. Energy conservation is the statement that

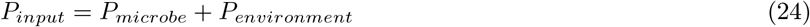

Plugging in the expression above yields the relationship

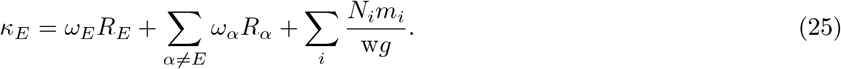

In the limit, where *P_microbe_* ≫ *P_environment_* this becomes

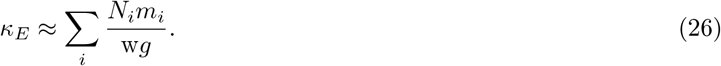

### Externally Supplied Resource

In our set-up, there is a single distinguished resource *R_E_* that is supplied externally. The dynamics of this resource are given by Eq. 12 with *κ_E_* ≠ 0. In fact, to ensure a good thermodynamic limit we must scale *κ_E_* with the number of resources *M* [13]. In other words, we have

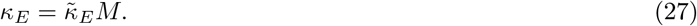

With this choice, we see that the steady-state abundance of the externally supplied resource will also scale extensively *R_E_* ~ *M*. For this reason, we can solve for the steady-state abundance of *R_E_* using “naive mean-field theory”. In particular, within naive MFT, the production and depletion flux in Eq. 12 can be replaced by their mean-field averages

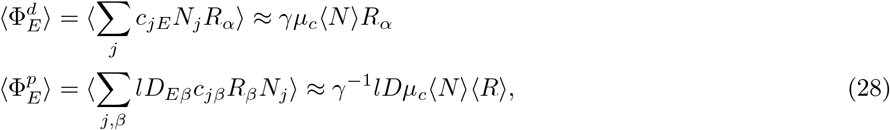

where we have defined the averages

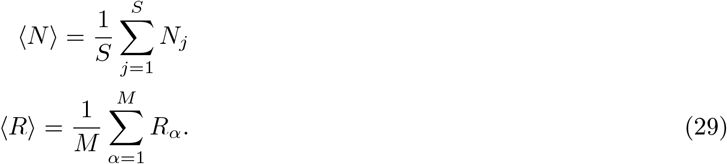

Plugging these equations into Eq. 12 and solving for the steady-state gives

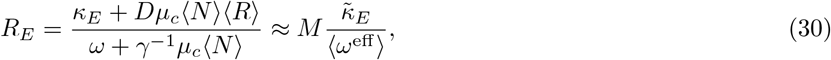

where in going to the second line we have dropped sub-leading order term in *M* and defined the average effective degradation rate

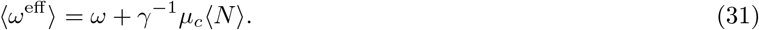

Thus, as expected the steady-state abundance of the externally supplied resource *R_E_* is extensive in *M*.

### Means, variances, and covariances for fluxes of self-generated resources

In the thermodynamic limit, we treat the production flux 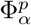 and depletion flux 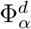, in Eq. 12 for self-generated resources as random variables. Under our assumption of *replica-symmetry*, such random variables are fully characterized by their means, variances, and covariances. To calculate these quantities, we make use of the cavity method.

In particular, we will consider a system in the *absence* of resource *α*. We will denote the steady-state abundances reached by species *N_j_* and resources *R_β_* in the absence of resource *α* by *N*_*j*/*α*_ and *R*_*β*/*α*_. Since through out this paper we will only be considering steady-states, our notation makes no distinction between steady-state quantities and dynamical quantities. We will then relate the means and variances of the fluxes 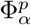 and 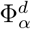, to *N*_*j*/*α*_ and *R*_*β*/*α*_ using perturbation theory. The logic behind this somewhat involved cavity method is that unlike naive-MFT this method properly accounts for all correlations in the problem.

### Depletion flux

Let us first write

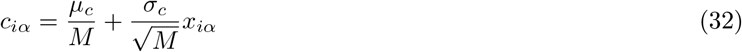

where the *x_iα_* are uncorrelated, standard normal variables. We first find

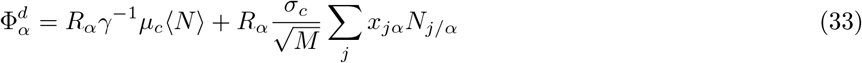

where through an abuse of notation we have defined the expectation value of 〈*N*〉 as

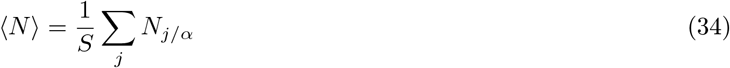

and *N*_*j*/*α*_ is the population of species *j* in the equilibrium state with resource *α* absent. This gives us that the mean of the depletion flux is just

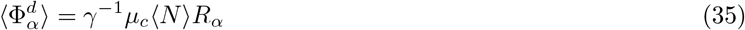

We can define

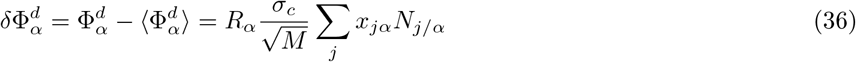

We can also calculate the variance of this flux

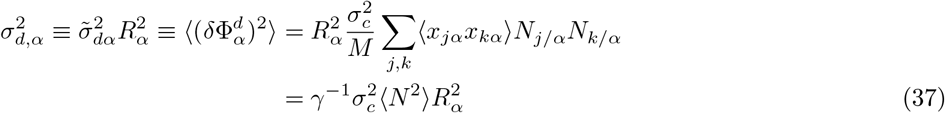

where we have defined

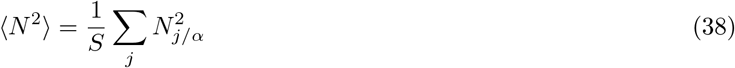

and have defined our shorthand symbol 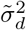 for this variance in a way that keeps the 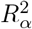 visible, since it will be important later on.

This allows us to approximate the production flux as a Gaussian random variable with

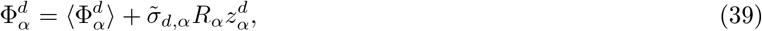

where 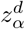 is a unit normal random variable and 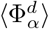 and 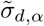 are defined in Eq. 35 and Eq. 37 respectively.

### Production flux

To calculate the mean and variance of the production flux, we once again start by making the dependence of *D_αβ_* of the *M* explicit by writing

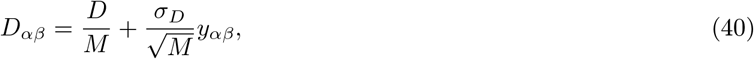

where the *y_αβ_* are uncorrelated, standard normal variables. This lack of correlation follows from the discussion of the Dirichlet distribution above. In terms of these variables we have

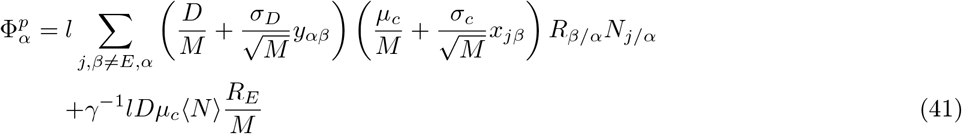

where once again we have treated the external resource in naive-MFT since it is extensive in *M*.

We have to take care in computing the mean flux, because there is an important correlations among *N_j_*, *x_jβ_* and *R_β_*:

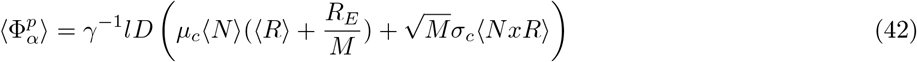

where in the last line we have used that fact this second term scales like 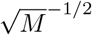 (see below for self-consistency proof) and we have defined

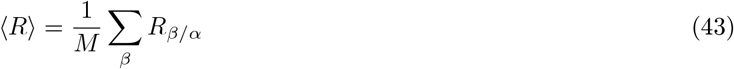

and

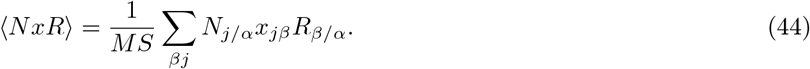

In writing this we have used the fact that *y_αβ_* is not correlated with *R*_*β*/*α*_ and *N_jα_* and hence all averages involving this quantity can be set to zero.

In addition to the mean value, to apply the cavity method we will also have to think about the fluctuations around the mean:

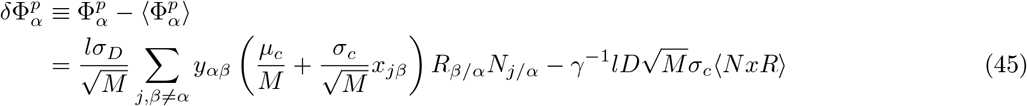

A straightforward but slightly tedious calculation shows that

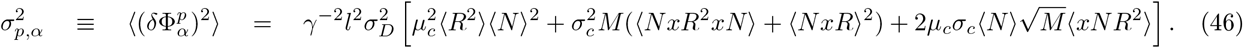

where again in the approximation we have ignored terms that scale as 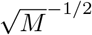 and defined

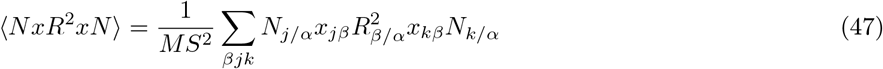

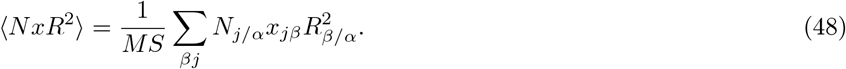

This allows us to approximate the production flux as a Gaussian random variable with

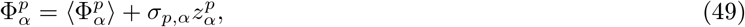

where 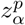 is a unit normal random variable and 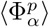 and *σ*_*p*,*α*_ are defined in Eq. 42 and Eq. 46 respectively.

### Covariance between production and depletion flux

From the above calculation, it is easy to show that to leading order in *M* that the covariance between the production and depletion flux is zero. To see this, we use Eq. 45 and insert this into the definition of the covariance 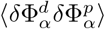 between the production and depletion flux. Since 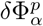 has a single factor of the standard normal variable *y_αβ_* multiplying all terms, the only way to get a nonzero average is for 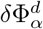 to contain factors of *y_αβ_* that could generate a 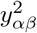. But the depletion flux does not have any dependence on *y_αβ_*, and so the correlation vanishes.

### Growth flux

In the previous section, we focused on the production and depletion fluxes for the resources. Here, we focus on the growth flux. Because we are using the cavity method, we consider a system where a species *i* has been removed from the ecosystem. The steady-state abundances reached by the ecosystem in the absence of species *i* are denoted by *N*_*j*/*i*_ and *R*_*α*/*i*_. We then ask about the growth flux that the species *i* would have when it invades such an ecosystem:

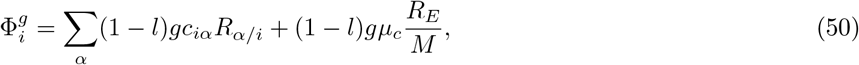

where once again we ignore fluctuations in the consumption of the externally supplied resource. As before, we write 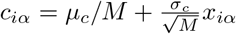 to get

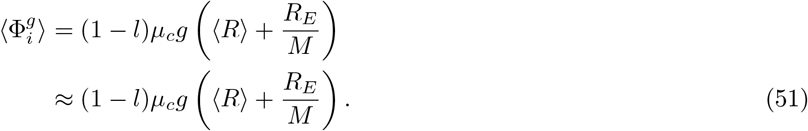

Defining the fluctuating part of the growth flux

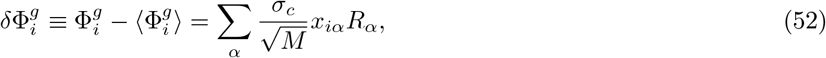

allows us to calculate the variance of the growth flux yielding:

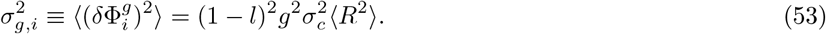

With these expressions, within the replica-symmetry ansatz the growth flux can be though of as a normal random variable of the form

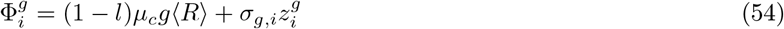

### Cavity Corrections

The essence of the cavity calculations is to relate an ecosystem with (*S* + 1, *M* + 1) species and resources to an ecosystem with (*S*, *M*) system. We will use the language of adding an additional species and resource which by convention we will denote by *N*_0_ and *R*_0_. The assumption is that when *S*, *M* ≫ 1, the addition of an additional species or resource is a small perturbation and we can relate the steady-state abundances in the presence and absence of the extra resource and species using perturbation theory.

### Setting-up the perturbation theory

In the presence of the new resource *R*_0_ and species *N*_0_, Eq. 12 is modified to as:

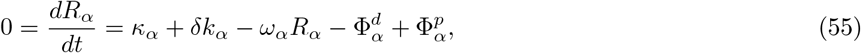

with

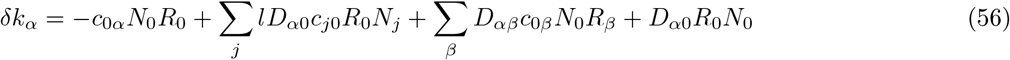

Similarly, Eq. 11 is modified to

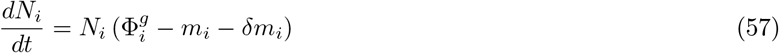

with

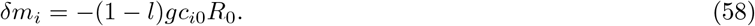

We can use perturbation theory to relate the species and resource abundances an ecosystem without the new resource and species (denoted by *N*_*i*/0_ and *R*_*α*/0_ respectively) to the abundances in the presence of *N*_0_ and *R*_0_ (denoted by *N_i_* and *R_α_* respectively). To do so, we define the susceptibilities

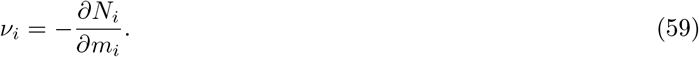

and

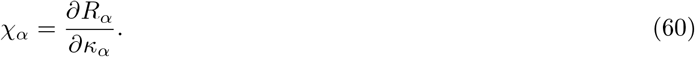

We note that this latter susceptibility differs from the resource susceptibility in [33] where we took the derivative with respect to *ω_α_* rather than *κ_α_*.

Combining the expressions above yields

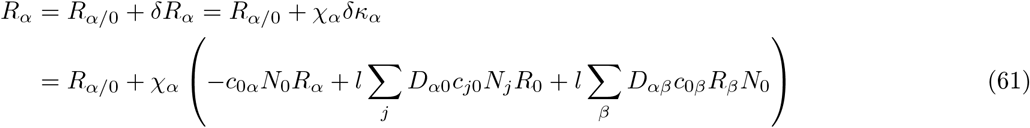

and

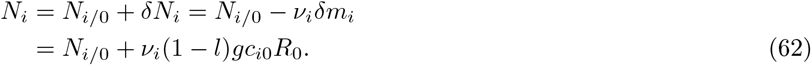

In writing these expressions, we have ignored the “off-diagonal” susceptibilities whose contributions we have assumed scale lower order in *M*.

### Computing the TAP corrections

To proceed, we need to calculate the TAP corrections to the three fluxes 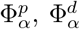, and 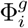 due to the addition of *N*_0_ and *R*_0_. Let us denote these cavity corrections to the fluxes by ΔΦ. Notice through an abuse of notation we will drop the /0 notation so that *R*_*α*/0_ is represented by *R_α_*. For the growth flux this amounts to calculating

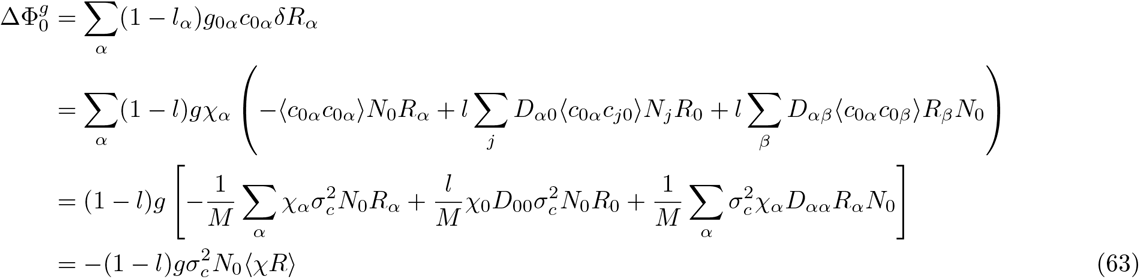

where

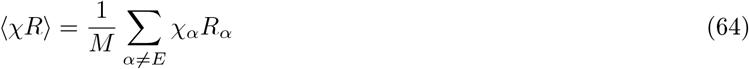

and we have dropped the last two terms from the third line because they are higher order in 1/*M*. We also need

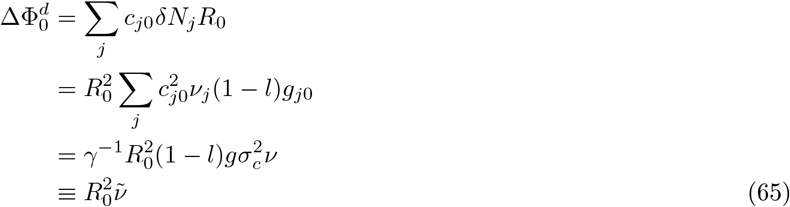

where we have introduced the notation

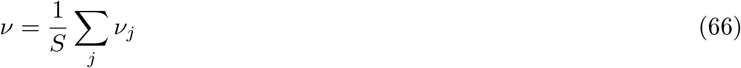

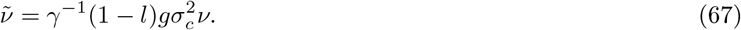

Finally,

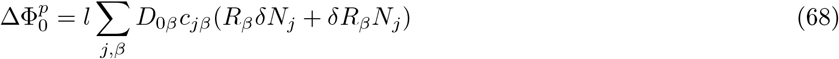

One can show that to leading order *M* that these terms disappear in the thermodynamic limit so that one has

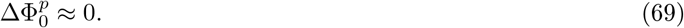

### Self-consistency equation

#### Equations for species

Let us start by defining an equation for species 0 at steady-state. We separate out the actual steady-state growth flux 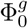 for species 0 into three parts: the average growth flux 〈Φ^*g*^〉 over all species in the regional pool, a Gaussian random variable with variance 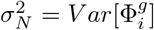 to account for the variability in growth rates in the regional pool, and finally the TAP correction 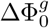 that accounts for the specific feedback from species 0 on the rest of the community:

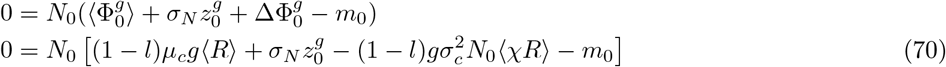

Requiring that the steady-states cannot be invaded (i.e are ecologically stable) translates into the statement that we choose *N*_0_ to be

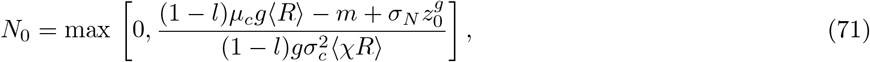

with

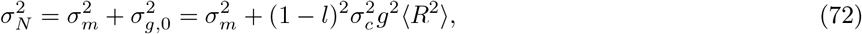

where in writing this equation we have used the fact that *m*_0_ is a normal random variable with variance *δm*^2^ that is uncorrelated with 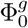 and Eq. 53.

Assuming replica symmetry, we can now write down the self-consistency equations for the moments of *N_i_*, using the special functions *w_n_* defined as

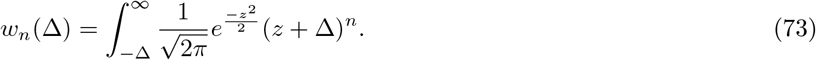

Using the usual arguments for replica-symmetric cavity solutions and matching the moments of Eq 71, one arrives at the expressions for the fraction of surviving species *ϕ_N_* and the first and second moments of the species abundance distributions 〈*N*〉 and 〈*N*^2^〉 [29–31]:

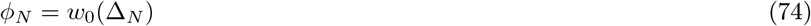

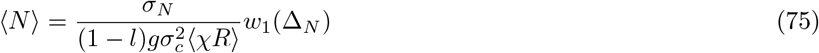

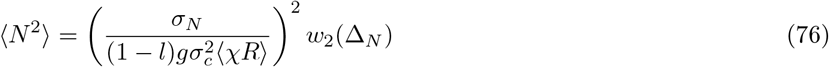

where

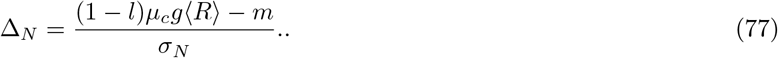

To calculate the average susceptibility we note that

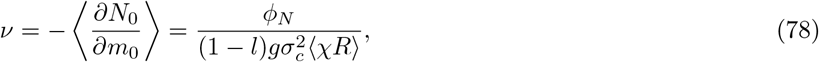

and, from Eq 67 that:

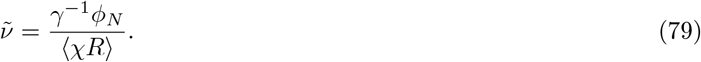

Eq. 71 also allows us to compute the expectation values that occur in Eq. 42 and Eq. 46. To do so, we note that from

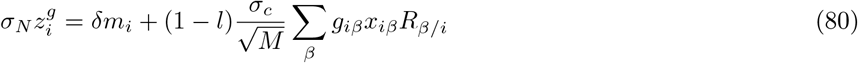

Using this expression and the definition of *x_iβ_*, we obtain:

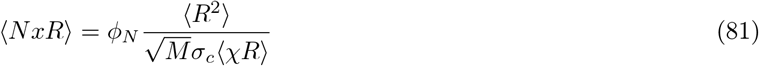

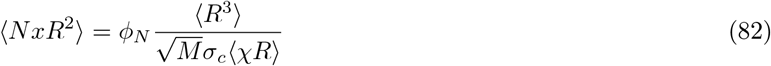

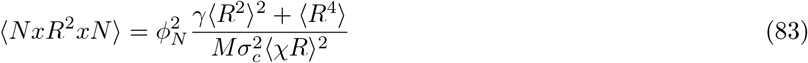

The first three quantities have the correct scaling for the production flux and its variance to remain finite as *M* → ∞. In calculating these expressions, we have ignored correlation between *x_iα_* and *R_α_* which we have assumed are higher order in 1/*M*. We check that this assumption is reasonable numerically.

With these expressions, we can rewrite Eqs. 42 and 46 as

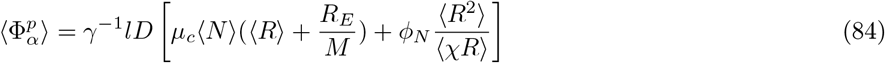

and

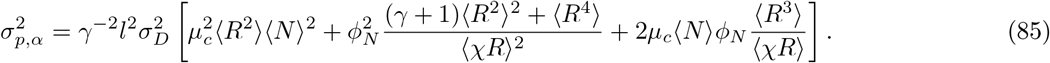

### Equations for resources

Let us now think about the steady-state equation for resource 0. We will separate the fluxes out into the same three components – the mean over the regional pool, a random term with variance equal to the variance over the regional pool, and a feedback term. Since we already showed that the covariance between the production and depletion fluxes vanishes to leading order in 1/*M*, we can introduce independent random terms for each of these fluxes. We recall that *κ*_0_ = 0 since we are considering a scenario with just a single externally supplied resource, but we will keep the term present for the sake of defining the susceptibilities. This gives:

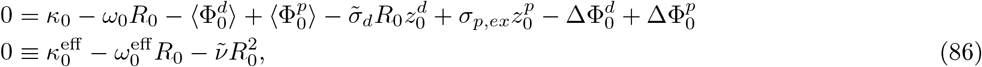

with

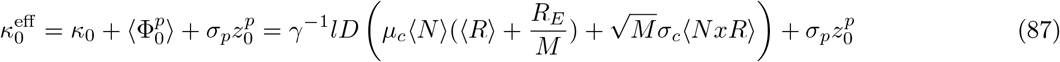

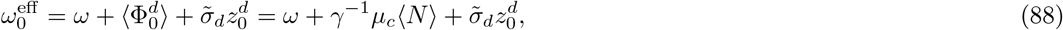

where in first line we have used *κ*_0_ = 0 and Eq. 42 and in the second equation we have used Eq. 35. This is identical to the case without cross-feeding considered in [33] with *κ* → *κ*^eff^, *ω* → *ω*^eff^

Solving for *R*_0_ gives:

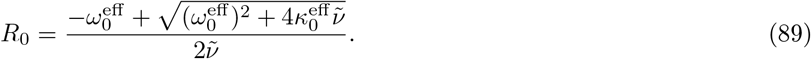

This expression allows us to compute self-consistency equations for the first two moments of the distribution of *R_α_*, which require performing some expansions in small parameters to solve, as in the case without crossfeeding. We can also compute

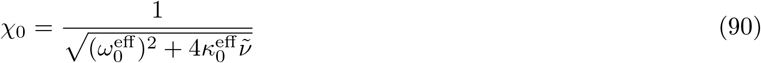

and find

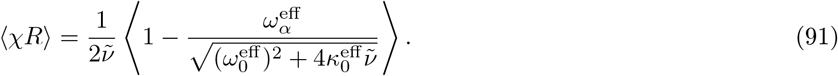

### Solving the equations for resources

Unfortunately, due to the non-linear nature of Eq. 89 it is not possible to solve for the moments of *R*_0_ without making additional approximations. In what follows, we will perform an expansion assuming the fluctuations in the production flux 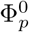 (Eqs. 42 and 46)and depletion flux 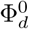 (Eqs. 35 and 37) are small. The first thing we need to do in order to solve these equations is to perform a Taylor expansion of the square root. To do so, we define three “dimensionless” parameters:

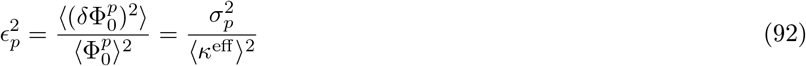

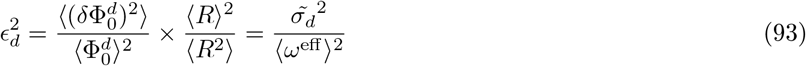

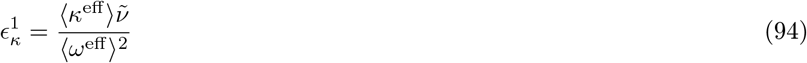

From these definitions, it is clear that the first two parameters measure fluctuations in the production and depletion flux respectively. The parameter *ϵ_k_* is a little harder to interpret. We will see below that the leading order contribution to 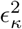 is the just the species-packing fraction and is always less than 1/2 (see derivation of species packing bound below).

In terms of these parameters, we have:

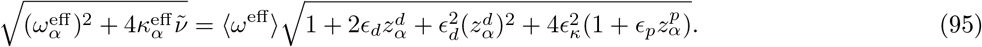

We will use the formula:

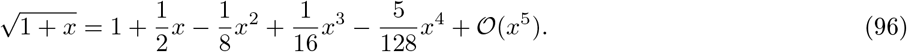

Keeping terms up to order *ϵ*^4^ (and noting that several terms cancel which greatly simplifies the final expression), we have

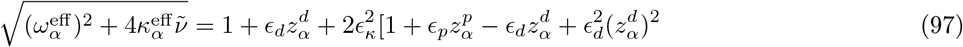

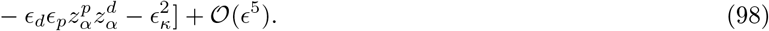

Using the expansion in eq. (98) in the expression for *R_α_* found in eq. (89) above, we have:

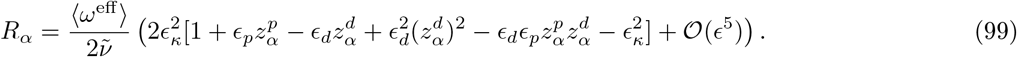

Now we can use the definition of *ϵ_κ_* to simplify the expression further:

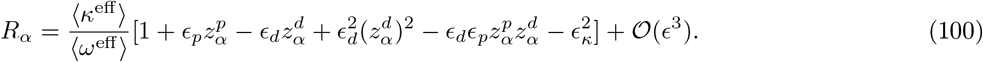

This expression makes sense, because to lowest order the resource abundance is just the ratio of supply to depletion. We can immediately calculate the expectation of various moments. The mean resource abundance is given by

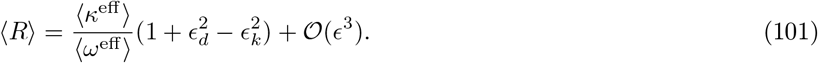

Similarly, one can express higher order moments as

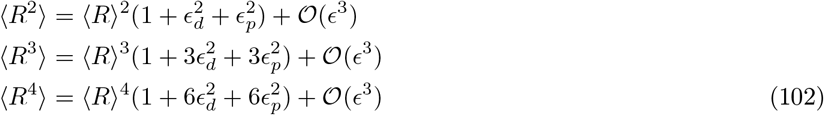

### Average abundance of leaked resources

We now solve for the average resource abundance 〈*R*〉. To proceed, we substitute the expression for 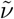 in Eq. 79 into Eq. 91 to obtain the intermediate relation

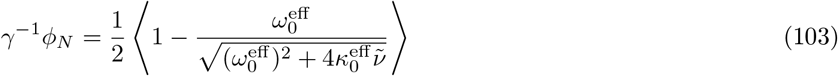

Using eq. (98), this can be rewritten as

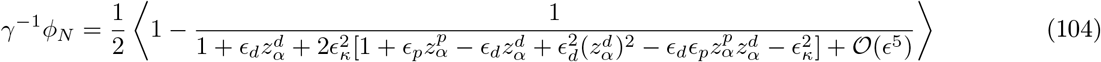

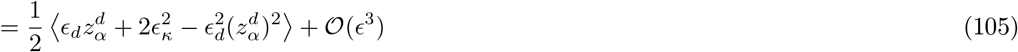

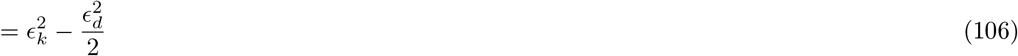

Explicitly this last equation can be written as

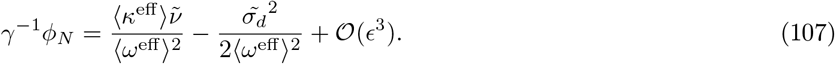

Rearranging this equation gives

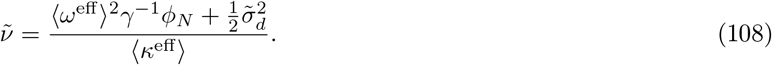

Using the expression for 〈*R*〉 in Eq. 100, we can simplify this to:

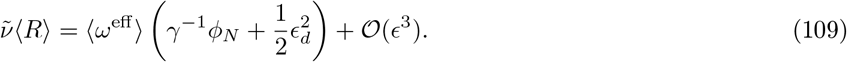

Combining this with Eq. 79 also yields the relation:

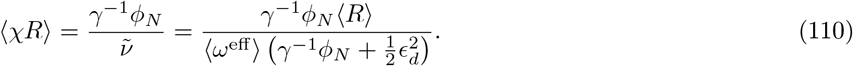

Substituting our expression for 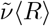, Eq. 83, and Eq. 101 into eq. (88) gives

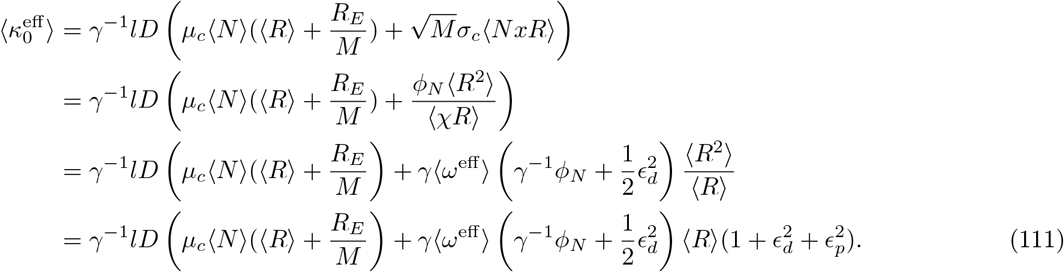

We can substitute this equation into Eq. 101 to get

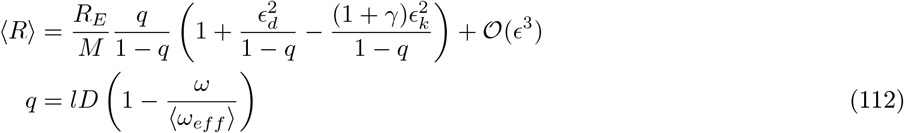

Substituting Eq. 30 into the expression above yields

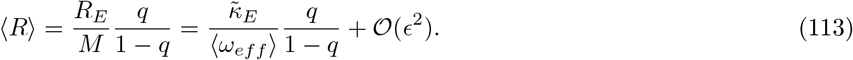

One final approximation we can make is consider the limit where depletion is dominated by microbes and we expand the expression above dropping all terms 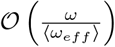. In this case, defining

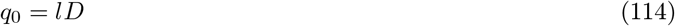

and using energy conservation relation Eq. 26 with w = 1 we have that

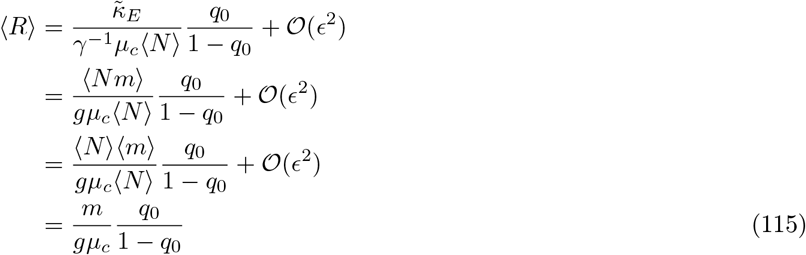

where we have defined the expectation

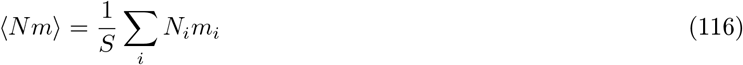

and in going from the second to third line we have used that 〈*Nm*〉 = 〈*N*〉〈*m*〉 at this order in *ϵ*.

### Second and higher-order moments of resource abundance

In order to calculate higher order moments of resources (see Eq. 102) we need to calculate 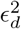 and 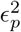. Substituting the expressions in Eq. 35 and Eq. 37 into Eq. 94 yields

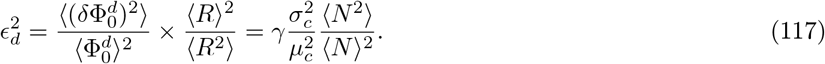

This expression implies that, as expected, 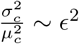.

In order to calculate 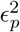, we start by calculating the average of the production flux. Combining Eq. 42 with Eq. 91 and yields Eq. 106 yields

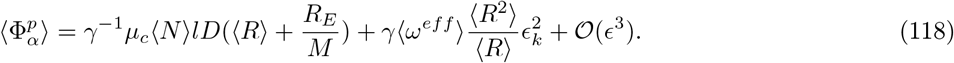

To simplify this expression further, we note that *γ*^−1^*μ_c_*〈*N*〉 = 〈*ω^eff^*〉 – *ω* and then use Eq 112 to get

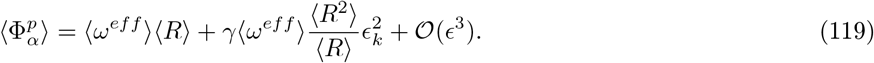

In order to calculate 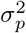 in Eq. 42, we start by simplifying the expressions in Eq. 83 using Eq. 110, Eq. 102, and Eq. 106, yielding:

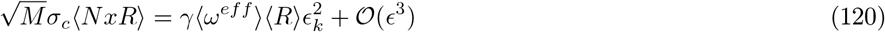

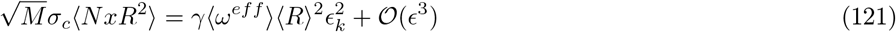

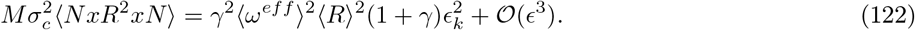

Plugging these into Eq. 42 gives

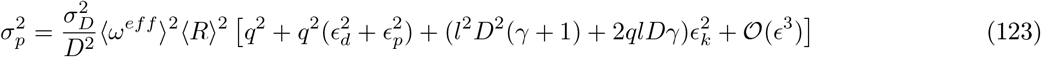

and Eq 46. This yields

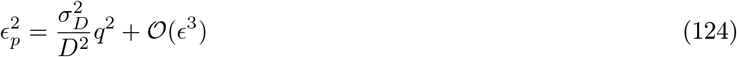

where we have noted this expression implies that as expected 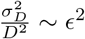. Thus, to this order in 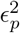 we have

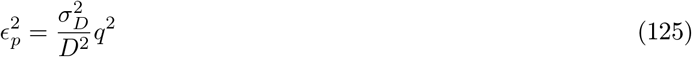

### Summary of results

In this section, we summarize our results from the calculation. We start by defining the parameters

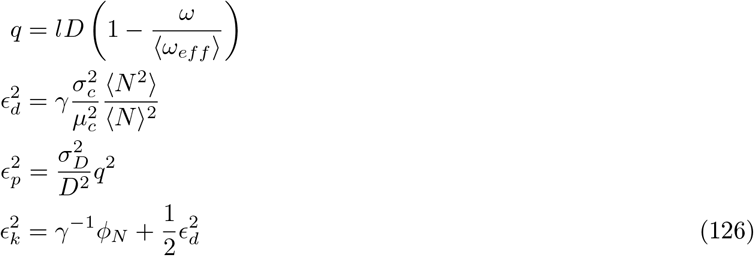

The average resource abundance is given by

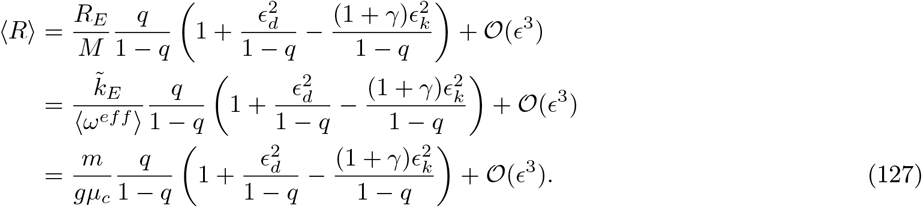

The higher order resource moments are just

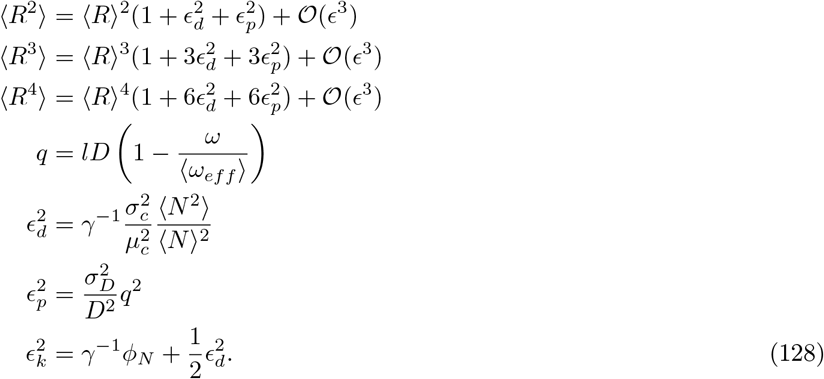

The full resource distribution is given by

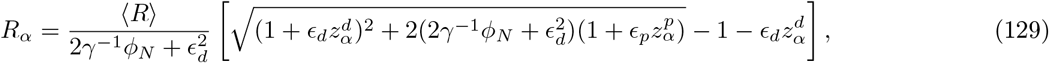

where are before 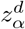 and 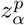 are standard random normal variables.

In addition, we have expressions for the species abundances.

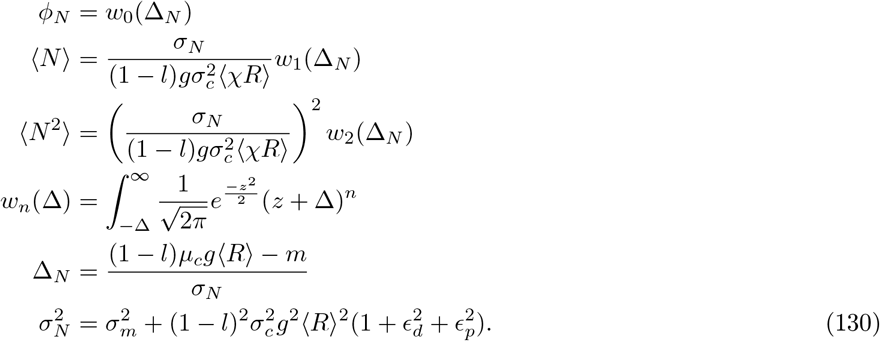

Finally, we have

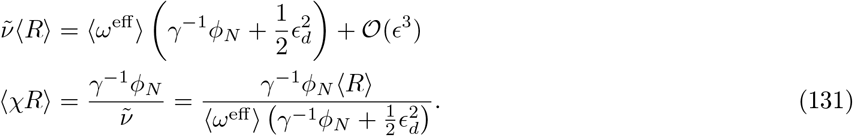

### Species Packing Bounds

The expressions above allow us to define a simple bound for the fraction of metabolic niches that an ecosystem can occupy [33]. If we denote the number of surviving species by *S*\* and the number of metabolites by *M*, then in the thermodynamic limit the species-packing fraction is just *S*\*/*M* = *γ*^−1^*ϕ_N_*. To derive the bound, we start with Eq. 91 which reads

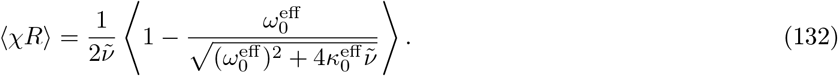

Substituting Eq. 79 for 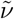 into the expression yields

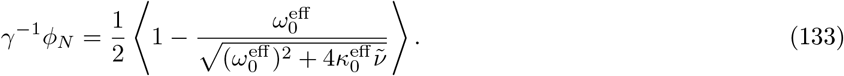

Since the second term is always positive we have that

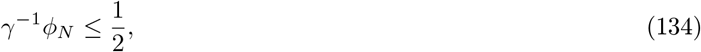

proving our bound. Note that in proving this bound, we have made no approximations beyond replica-symmetry and therefore expect this to be exact in the thermodynamic limit.

### Fraction of surviving species depends only on competition and not leakage rate

In this section, we show that to leading order in *ϵ* (see Eq. 94), the fraction of surviving species depends only on the competition between species (*μ_c_* and *σ_c_*) but not the leakage rate *l* when 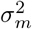, the fluctuations in the maintenance costs, can be ignored. Technically, this is a statement that Δ_*N*_, and hence *ϕ_N_*, depends on the variance over the mean squared of consumer preferences 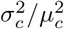 but not the leakage parameter *l*. There are many ways to show this and here we present one particularly simple derivation of this fact.

Our starting point is Eq. 130:

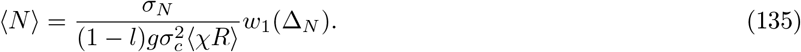

We then use Eq. eq:chiR2 to rewrite this as

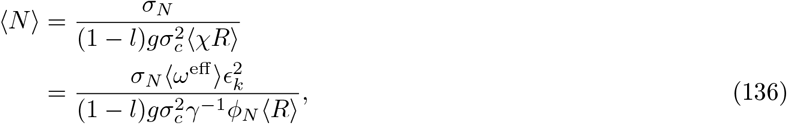

where in the second line we have used Eq. 110 and Eq. 126 for 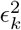. Now, using the fact that when *σ*^2^ =0 we have from Eq. 130 that

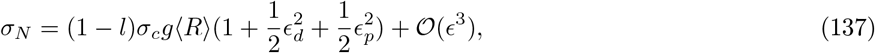

the above can be rewritten as

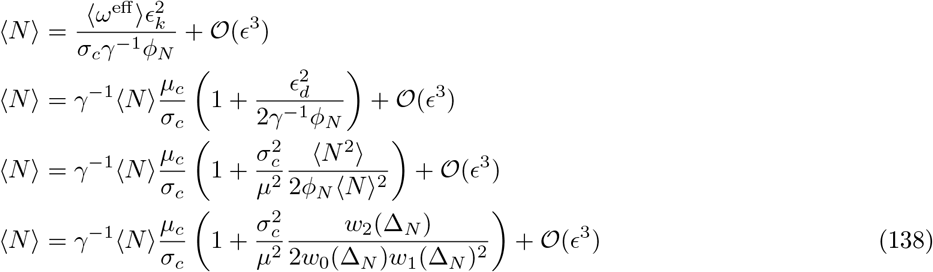

Canceling the 〈*N*〉 from both sides yields

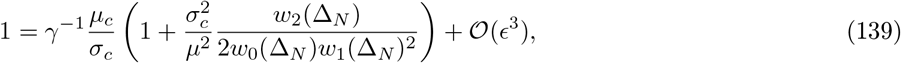

showing that Δ_*N*_ and hence *ϕ_N_* only depends on 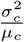 at this order.

### Details of numerical simulations

#### Numerical Simulations

We now give an overview of how we performed numerical simulations. All code can be found on the corresponding Github page. To numerically simulate the communities, we used the Community Simulator package to find steady-states with indicated parameters [21]. We used the parameters: *q* = 0 (generalist consumers), supply = ‘external’, regulation= ‘independent’, response=‘type I’, and sparsity *s* = 0.01. These choices ensure that the dynamics follow Eq. 12. We set the number of species *S* = 600, the number of metabolites equal *M* = 300, the average maintenance cost *m* = 1 with standard deviation *σ_m_* = 0.1. A single resource was supplied externally at a rate 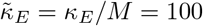 and *ω* = 1. The leakage rate was chosen to be *l* = 0.8 if fixed, or varied over 10 equal intervals from *l* = 0.7 – 0.95 as indicated in figures.

The consumption coefficients were chose from binary distribution with *c_iα_* ℰ {0, *c*_1_} with probability having nonzero coefficient *c*_1_ given by *p* = *μ*_*c*_/(*c*_1_ *M*). This was done to ensure that the average consumption rate 〈*c_iα_*〉 = *μ_c_*/*M*, with *μ_c_* = 1. Under these assumptions 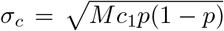. In order to vary competition, *c*_1_ was varied over 10 equal interval from *c*_1_ = 1/*M* – 10/*M* with *M* = 300. Default choice for *c*_1_ = 9/*M* = 0.03. See section “Sampling parameters” in [21] for details and Github for code.

Averages *ϕ_N_*, 〈*N*〉, 〈*N*^2^〉, 〈*R*〉, and 〈*R*^2^〉 reflect averages over 28 independent realizations. In order to ensure numerical stability, we treat species as abundances below 10^−3^ as going extinct.

### Numerical solution of cavity equations

To numerically check the solutions to our self-consistent cavity equations, we used the expressions summarized in section. In particular, given the parameters we solved for Δ_*N*_ numerically. To do so, we used the second line of Eq. 138, Eq. 128, and Eq. 130 to get:

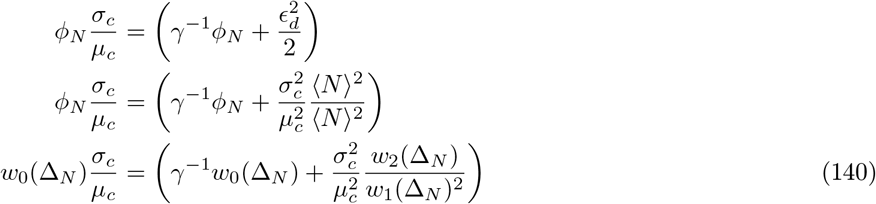

This allowed us define the cost function

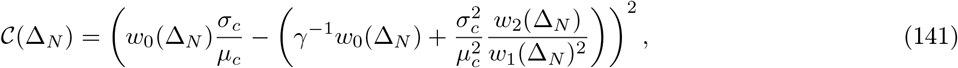

which we can minimize to solve for Δ_*N*_ using the “minimize scalar” function from the scipy.opitmize package.

Once we solve for Δ_*N*_, we can use Eq. 126 along with Eq. 129 to numerically calculate the predicted resource distribution:

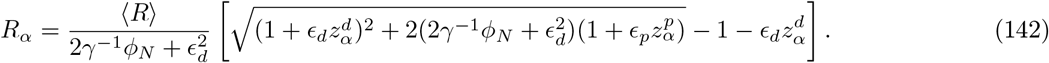

To do so, we randomly sample from this distribution 10^6^ times and then plot the resulting pdf.

We can also use Eq. 71 to generate the distribution of the species abundances. In particular, this equation states that the distribution of *surviving species* follows a truncated Gaussian distribution with a fraction *ϕ_N_* of the species surviving. To proceed, note that from Eq. 130 that we can define the quantity

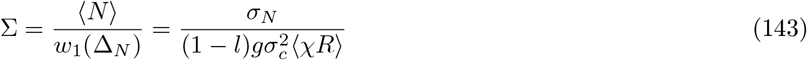

In terms of Σ, we can rearrange Eq. 71 to derive that *surviving species* will be described by the distribution

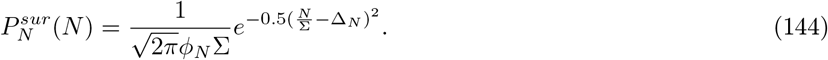

Δ_*N*_ is obtained by minimizing 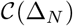 as discussed above. To obtain 〈*N*〉 to leading order in *ϵ*, we use energy conservation Eq. Eq:EnergyCons:

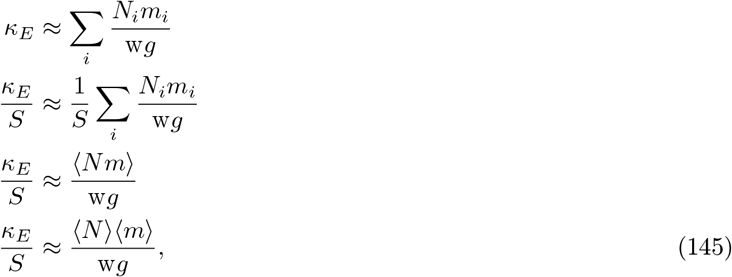

which can be rearrange to yield

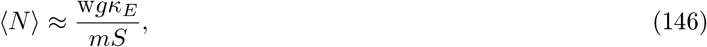

allowing us to calculate Σ.

As a further check the prediction of the cavity method and our approximations we use the value of Δ_*N*_ obtained from the procedure above along with Eq. 130 to compute

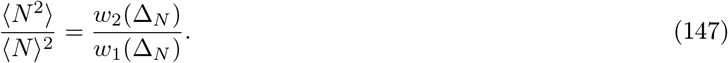

**FIG. S1.**
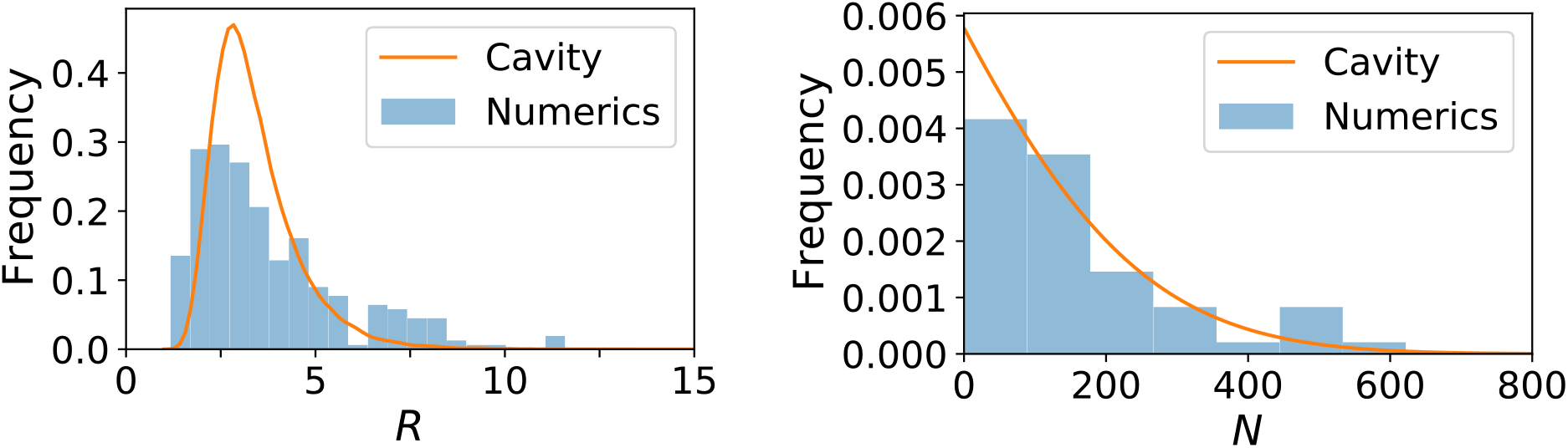
Cavity solutions predict distributions of species and abundances. The resource and species abundance distributions from numerics and our cavity solutions Eqs. 7 and 5. See appendix for parameters.

To check that the average resource abundance 〈*R*〉 only depend on the leakage rate in the limit of fast dilution (i.e. when *q* ≈ *q*_0_ = *l*), we plotted the leading order contribution to Eq. 127:

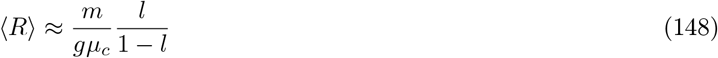

versus the average obtained from direct numerical simulation.

## Additional Figures and Numerical Checks

**FIG. S2.**
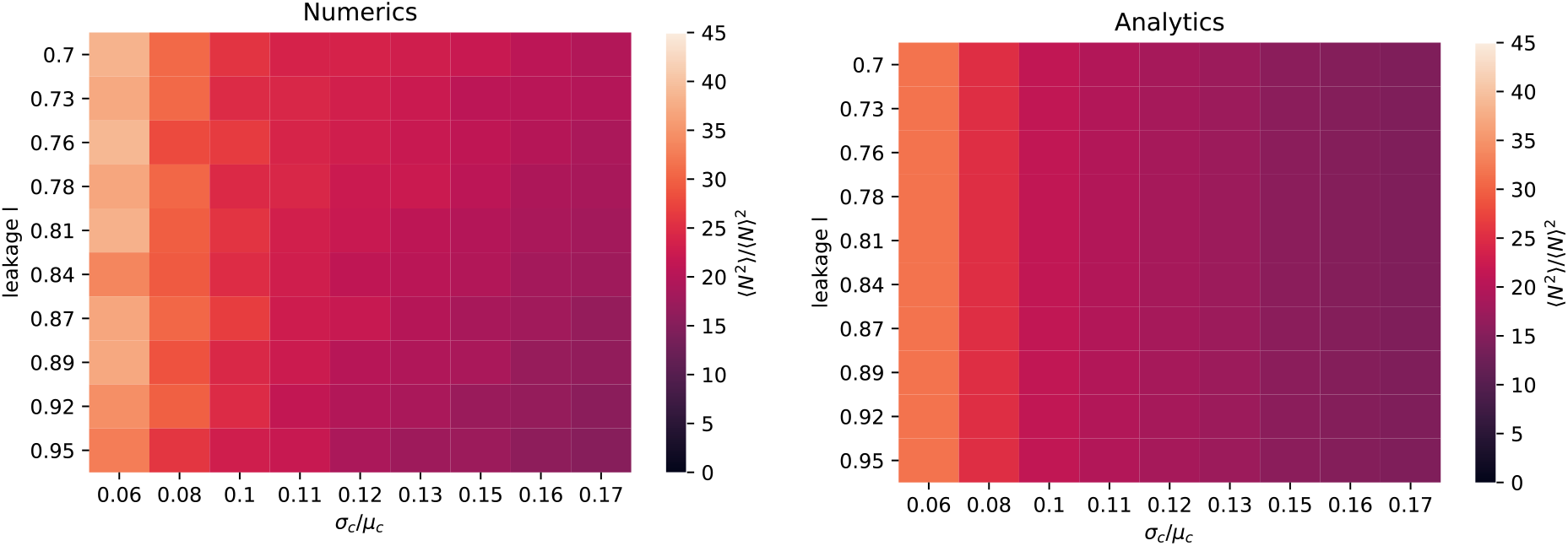
Species distributions set by competition. Numerical simulations (left) and analytics (right) of 〈*N*^2^〉/〈*N*〉^2^ as a function of leakage rate *l* and *σ_c_*/*μ_c_* given by Eq. 138 for binary consumer resource coefficients as described above.

**FIG. S3.**
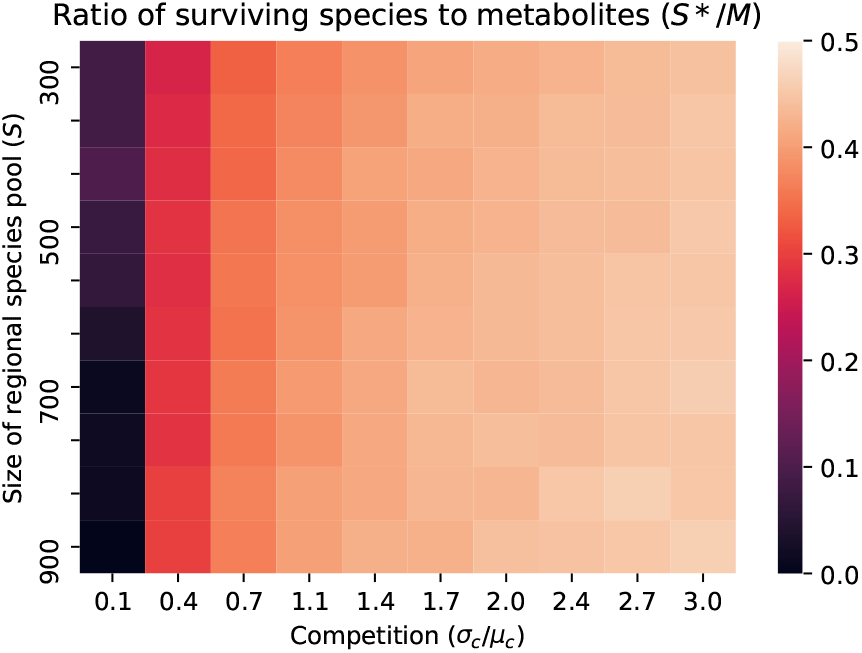
Species packing bound with Gaussian consumer preferences. Ratio of surviving species to resources (including metabolic biproducts) *S*\* /*M* for different leakage rates *l* and different choices of *σ_c_*/*μ_c_* with size of regional species pool *S* = 600 and *M* = 300. Notice that *S*\*/*M* ≤ 1/2, consistent analytic bound derived in appendix.

